# A role for cerebral cortex in the suppression of innate defensive behavior

**DOI:** 10.1101/2020.12.22.424023

**Authors:** Silvia Natale, Maria Esteban Masferrer, Senthilkumar Deivasigamani, Cornelius T. Gross

## Abstract

The cerebral cortex is involved in the control of cognition and the processing of learned information and it appears to have a role in the adaptation of behavior in response to unpredictable circumstances. In addition, the cortex may have a role in the regulation of innate responses since rodents, cats or primates with surgical removal or accidental destruction of cortical regions show excessive irritability, aggression and rage elicited by threatening stimuli. However, it remains unclear whether cortex has an acute role in suppressing innate threat responses because the imprecision and chronic nature of these lesions leaves open the possibility that compensatory processes may underlie some of these phenotypes. In the present study we used pharmacogenetic inhibition to precisely, rapidly and reversibly suppress cortical pyramidal neuron function and examine its contribution to defensive behaviors elicited by a variety of innately aversive stimuli. Inhibition of cortex caused an increase of defensive responses elicited by an aggressive conspecific, a novel prey, and a physically stressful stimulus. These findings are consistent with a role of cortex in the acute inhibition of innate defensive behaviors.

## Introduction

The cerebral cortex is an evolutionarily recent brain region that comprises the largest part of the mammalian forebrain (Molnár et al., 2014; Montiel and Aboitiz, 2015; Merel et al., 2019). Lesions of the cortex reduce cognitive and motor function in primates, but in lower mammals the motor effects of such lesions are minimal (Walker and Fulton, 1938; Lopes et al., 2017), suggesting that the executive motor control function of the cerebral cortex is a relatively recent adaptation. This distinction is supported by anatomical evidence that cortical projection neurons directly enervate motor neurons in spinal cord only in primates (Bernhard & Bohm, 1954; Lawrence & Kuypers, 1968; Yang & Lemon, 2003). As a result of the more severe effects of cortical lesions in higher mammals it has been argued that the primary function of the cerebral cortex is better understood by examining its contributions to brain function in lower mammals (Lopes et al., 2017). Cortical lesions in cats, for example, spare motor function and simple learning, although learning is slower and the animals show hesitation in executing motor actions that require three dimensional visual-motor feedback (Bjursten et al., 1976). Decorticated animals eat and drink adequately, show normal sexual behavior, and maintain normal locomotor activity (Bjursten et al., 1976). They also perform visual discrimination (Winans, 1967) and are able to retain memories acquired before the lesion (Kawai et al., 2015). These data are consistent with the hypothesis that the primary role of cerebral cortex is to acquire experience-dependent models of the world and use these to guide behavior in moments of uncertainty (Marple-Horvat et al., 1993; van Bergen et al., 2015; Heindorf et al., 2018; Schröder et al., 2019).

In contrast, subcortical brain regions are evolutionarily ancient, having their origins in the annelid and possibly pre-bilaterian ancestor (Tessmar-Raible et al., 2007). Notably, the hypothalamus contains a series of motor control networks that are necessary for the formation of internal states and the production of stimulus-appropriate adaptive behavioral and physiological responses, in particular those concerned with reproduction, defense, and seeking (Swanson, 2000). Thus, the mammalian brain is able to elicit the majority of essential behavioral responses using subcortical circuits, while the cerebral cortex is likely to have developed to provide cognitive support for motor actions, culminating in the development of its ability to directly control motor behaviors in primates (Tomer et al., 2010).

Nevertheless, early lesion studies noted that removal of the cerebral cortex altered more than just cognitive function. In the first case in which such a lesion was carefully documented, damage to the human frontal cortex was associated with irritability, irreverence, and disrespectfulness, with the attending physician noting that the “equilibrium between intellectual faculties and animal propensity had been destroyed” (Harlow, 1869). Later, similar phenotypes were found in laboratory cats and dogs with targeted lesions of the cerebral cortex – that consistently showed exaggerated defensive responses to otherwise innocuous stimuli (Goltz, 1892; De Barenne, 1920; Rothmann, 1923). Further studies showed that the phenotype extended to hyperexcitability caused by noise or by a physically stressful condition such as infection by worms (Schaltenbrand and Cobb, 1931). These so called “sham rage” phenomena were hypothesized to derive from the release from cortical inhibition of midbrain regions known to generate defensive behavior (Cannon and Britton, 1925; Bard,1928; Bard, 1934; Bard, 1937). Interestingly, when the decortication was performed during infancy, lesioned animals only started to show exaggerated defensive responses following sexual maturity (Bjursten et al., 1976) suggesting that this cortical inhibition function develops during adolescence.

However, cortical lesion studies to date have used physical removal of the cortical hemispheres and it remains possible that the effects seen depend on long-term compensations that may have occurred in the remaining brain regions. At the same time, aspiration lesions of the cortex frequently suffer from imprecision that spares parts of cortex or removes adjacent sub-cortical regions (Goltz, 1881, 1888) leaving it unclear if the lesion phenotypes were strictly dependent on cortical function (Head, 1921). Finally, the impact of cortical lesions on innate behaviors has not been systematically examined. In particular, it remains unclear whether the increased threat responses seen in some studies generalize across stimuli. To overcome these limitations and directly assess whether cortex has a generalized role in the acute inhibition of threat responses we developed a pharmacogenetic method to precisely, rapidly, and reversibly suppress cortical function in mice and assess alterations in a variety of innate defensive behaviors. We found that acute inhibition of cortex caused increased defensive responses to social, non-social, as well as physical threat cues.

## Results

### Rapid pharmacogenetic inhibition of cerebral cortex

In order to precisely, rapidly, and reversibly inhibit the entire cerebral cortex we generated mice expressing the pharmacogenetic inhibitory receptor hM4D (Armbruster et al., 2007) in all cortical principal neurons. Experimental and control littermate mice were obtained by crossing mice expressing hM4D under a ubiquitous Cre-dependent promoter (*RC*::PDi, Ray et al., 2011) with mice expressing Cre recombinase under control of the *Emx1* promoter (*Rosa*-*CAG*::LoxP-mCherry-STOP-LoxP-hM4D x *Emx1*::*Cre*; **Figure 1A**). *Emx1*::*Cre* drives expression exclusively in excitatory projection neurons of the cerebral cortex. Importantly, the driver line is active in both neocortex and archicortex, and thus allowed us to precisely direct hM4D expression in the entire cerebral cortex in a manner not possible with earlier lesions (Iwasato et al., 2000; Iwasato et al., 2004). Control animals lacked the *Emx1*::*Cre* allele and thus did not express hM4D (**Figure 1A**). As expected, fluorescence of the mCherry reporter was found throughout the brain in control mice (**Figure 1B**), but was reduced in the cortex of experimental, double transgenic animals (**Figure 1C**). Immunofluorescence staining against the hemaglutinin (HA) tag of hM4D showed expression in cortex of experimental, but not control animals, while expression in thalamus was lacking in both, as expected (**Figure 1BC**).

**Figure 1.**
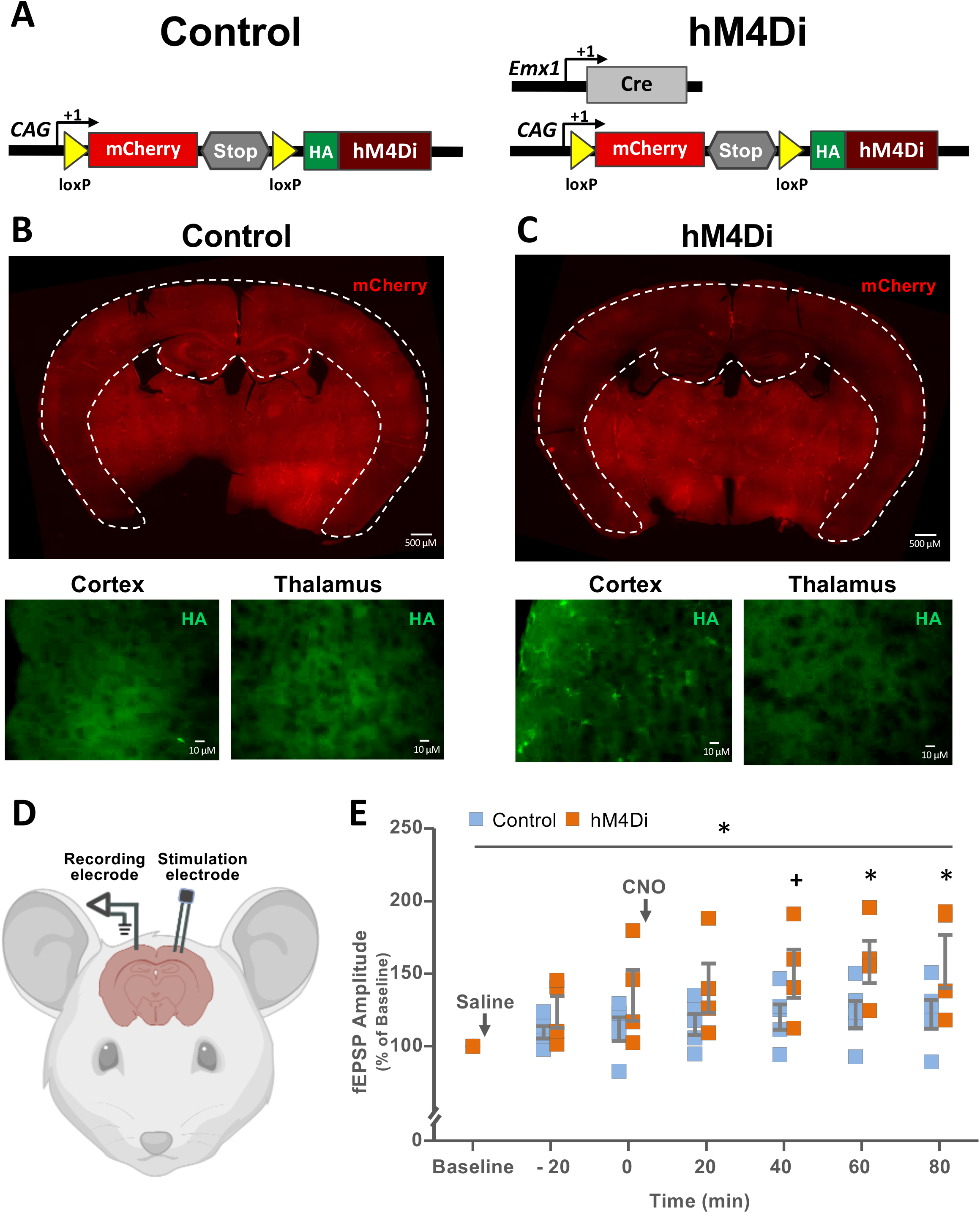
Transgenic mouse allowing rapid drug-induced cortical inhibition. **(A)** Schematic representation of (**Left**) Control and (**Right**) hM4Di experimental mice carrying, respectively, the Cre-dependent transgene expressing the pharmacogenetic inhibitory receptor hM4Di alone or together with the tissue and cell-type specific Cre drive line *Emx1*::Cre. In the absence of Cre activity, mCherry is ubiquitously expressed and hM4Di is not expressed, while in the presence of Cre activity, mCherry is excised and hM4Di is expressed. **(B)** Histological analysis demonstrating (**top**) ubiquitous mCherry fluorescent reporter gene expression and background staining for the hemagglutinin-tagged hM4Di receptor in (**bottom left**) cortex and (**bottom right**) thalamus. **(C)** Histological analysis demonstrating (**top**) selective deletion of mCherry fluorescent reporter gene expression in neocortical and archicortical structures and positive staining for the hemagglutinin-tagged hM4Di receptor in (**bottom left**) cortex, but not (**bottom right**) thalamus. **(D)** Schematic representation of *in vivo* electrophysiology recording set up showing stimulating and recording electrodes implanted contralaterally in the cortex. (**E**) Quantification of field excitatory postsynaptic potentials (fEPSPs) recorded in Control (N = 5) and hM4Di (N = 4) mice at various times before and after CNO treatment (^+^P < 0.1, *P < 0.05, **P < 0.01, ***P < 0.001).

Treatment of mice expressing hM4D with clozapine-N-oxide (CNO) results in the activation of hM4D and the activation or inhibition, respectively, of somatodendritic G-coupled inward rectifying potassium (GIRK) and presynaptic voltage-gated calcium (VGCC) channels and inhibition of neural responsivity and neurotransmitter release (Armbruster et al., 2007; Mizutani et al., 2006). To confirm neural inhibition of cortex in our animals we carried out *in vivo* electrophysiological measurements of evoked local field potential (fEPSP) responses in experimental and control littermates treated with CNO. Recording and stimulation electrodes were implanted contralaterally in the motor cortex (Oh et al., 2014; **Figure 1D**) and fEPSP responses to repeated stimulation (0.1 Hz, 1-10 mA) were collected and analyzed. Consistent with previous data in which fEPSPs were monitored to assess hM4D function (Madroñal et al., 2016) we observed a significant difference in fEPSP amplitude in experimental versus control animals starting 1 hour after CNO delivery (3 mg/kg, i.p.; two-way Repeated Measures ANOVA – treatment: F[1, 42] = 2.93, P = 0.131; Time: F[6, 42] = 14.29, P < 0.001; treatment x time: F[6, 42] = 2.78, P = 0.023; **Figure 1E)**. We noted that fEPSP amplitude was increased by CNO treatment in hM4D expressing mice compared to control animals, a phenomena consistent with the expression of post-inhibitory excitation following GPCR-dependent hyperpolarization of cortical pyramidal neurons (Spain et al., 1991; Grenier et al., 1998). These data demonstrate a rapid inhibition of cortical excitatory neurotransmission in our experimental animals.

### Cerebral cortex suppresses social avoidance

To examine a possible role of cerebral cortex in the modulation of innate defensive responses we treated experimental mice with CNO to suppress cortical neuron function and exposed them to three classes of innate threat stimuli, a social aggressor, a novel prey, and a physical stressor. For exposure to social threat singly housed male hM4D and control mice were injected with saline or CNO (3 mg/kg, i.p.) and allowed to habituate to the testing room in their home cage for 1 hour before behavioral testing. Testing comprised a five minutes habituation phase followed by a five minutes social phase when a CD1 aggressor was placed into the home cage of the mouse within a wire mesh cage (Franklin et al., 2017; **Figure 2A**). Quantification of behavior showed that CNO-treated hM4D mice showed a significant decrease in total activity during the social phase when compared to saline-treated hM4D mice (two-way ANOVA – genotype: F[1, 26] = 0.451, P = 0.508; treatment: F[1, 26] = 2.63; P = 0.117; genotype x treatment: F[1, 26] = 10.31; P = 0.003), but that this effect was not present during the habituation phase (two-way ANOVA – genotype: F[1, 26] = 1.55; P = 0.223; treatment: F[1, 26] = 1.64; P = 0.211; genotype x treatment: F[1, 26] = 0.301; P = 0.588; **Figure 2BC**). CNO-treated hM4D mice also showed a significant decrease in social approach (two-way ANOVA – genotype: F[1, 26] = 0.317, P = 0.578; treatment: F[1, 26] = 1.72, P = 0.201; genotype x treatment: F[1, 26] = 4.98, P = 0.034) and increase in latency to investigate the threat during the social phase when compared to saline-treated hM4D mice (Mann-Whitney: U = 6.0, P = 0.017; **Figure 2DE**). No significant modulation of latency to (Mann-Whitney: U = 31.0, P = 0.959) or time spent in social approach (two-way ANOVA – genotype: F[1, 26] = 0.317, P = 0.578; treatment: F[1, 26] = 1.72, P = 0.201; genotype x treatment: F[1, 26] = 4.98, P = 0.034) or activity in the social (two-way ANOVA – Genotype: F[1, 26] = 0.451, P = 0.508; treatment: F[1, 26] = 2.63, P = 0.117; genotype x treatment: F[1, 26] = 10.31, P = 0.003) or habituation (two-way ANOVA, Genotype: F[1, 26] = 1.55, P = 0.223; treatment: F[1, 26] = 1.64, P = 0.211; genotype x treatment: F[1, 26] = 0.301, P = 0.588) phases was observed in control mice treated with CNO compared to saline, confirming that the effect of CNO in hM4D mice depends on the expression of the pharmacogenetic transgene (**Figure 2B-E**). Importantly, CNO-treated hM4D female mice exposed to the CD1 male did not show any change in activity (two-way ANOVA – genotype: F[1, 25] = 0.150, P = 0.702; treatment: F[1, 25] = 1.089, P = 0.307; genotype x treatment: F[1, 25] = 0.0461, P = 0.832) or social approach (two-way ANOVA – genotype: F[1, 25] = 0.0610, P = 0.807; treatment: F[1, 25] = 0.279, P = 0.602; genotype x treatment: F[1, 25] = 0.0168, P = 0.898) behavior when compared to saline-treated hM4D littermates, demonstrating that the effect of cortical inhibition is absent when the social interaction is affiliative, rather than antagonistic (**Figure S1B-D**). Nevertheless, a significant increase in latency to investigate in CNO-treated hM4D female mice when compared to saline-treated hM4D littermates supports the idea that the cortex is involved in the inhibition of avoidance when the threat is unknown (Mann-Whitney: U = 10.0, P = 0.021; **Figure S1E**).

**Figure 2.**
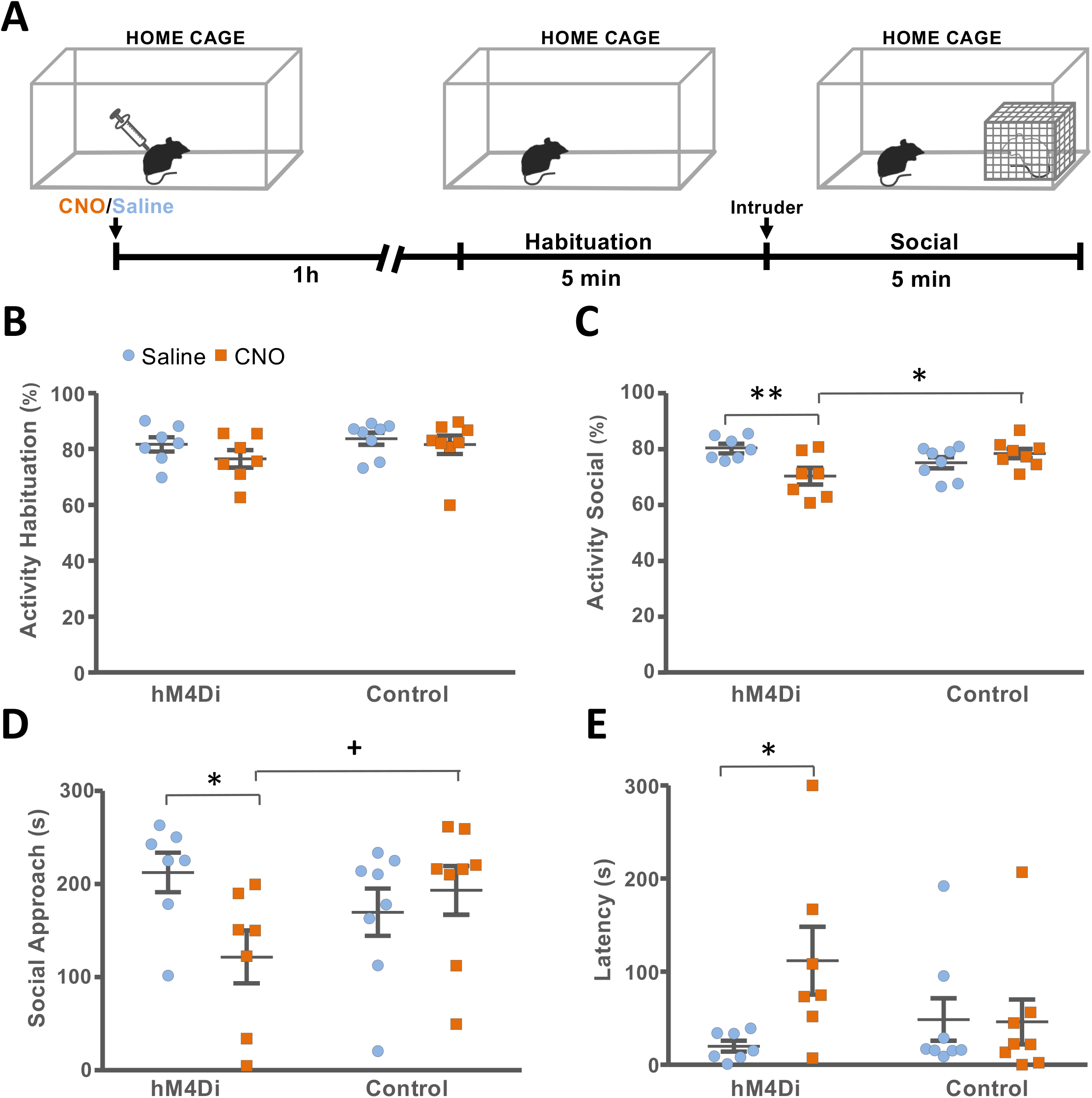
Acute cortical inhibition increases social avoidance. (**A**) Experimental protocol for testing social avoidance. Mice were treated with saline or CNO and returned to their home cage for one hour before baseline testing for five minutes followed by the introduction at one side of the home cage of an aggressor confined to a wire mesh cage and further observation for five minutes. Activity of mice measured during (**A**) habituation and (**B**) social test phases. (**C**) Time spent in social approach and (**D**) latency to first social approach (N = 7-8, ^+^P < 0.1, *P < 0.05, **P < 0.01, ***P < 0.001).

Next, we examined the effect of cortical inhibition on social approach in animals that had been habituated to the threat. Habituation reduces novelty-associated defensive behaviors and if cortex were to have a selective effect on defense, then the effect of cortical inhibition should be attenuated under these conditions. The social approach test was repeated daily for five consecutive days with CNO or saline treatment on the first and last days, under a pseudorandomized assignment (**Figure S2A**). To control for possible effects of stress associated with the treatment all animals were give saline on the intervening days. On day five, CNO-treated hM4D mice showed a significant increase in social approach as compared with the first day, reaching the same level as controls (two-way ANOVA – genotype: F[1, 23] = 1.31, P = 0.264; treatment: F[1, 23] = 6.069, P = 0.022; genotype x treatment: F[1, 23] = 4.68, P = 0.041; **Figure S2B**). As expected CNO treatment had no effect on total activity on social phase of either day (two-way ANOVA – genotype: F[1, 27] = 0.496, P = 0.487; treatment: F[1, 27] = 2.104, P = 0.158; genotype x treatment: F[1, 27] = 0.0229, P = 0.881; **Figure S2C**). This finding demonstrates that cortical inhibition selectively affects defensive behavior under conditions of novelty, and also supports earlier findings showing that habituation learning is intact in decorticated animals (Kawai et al., 2015).

Finally, we performed a within subject analysis to examine whether cortical inhibition selectively modulated defensive behaviors in a manner that was time-locked to the introduction of the threat stimulus. Individual and average plots of continuous freezing (**Figure 3A**) and avoidance (**Figure 3B**) behavior during the habituation and social phases revealed that CNO treatment was associated with a significant increase in freezing (two-way repeated measures ANOVA – treatment: F[1, 12] = 4.104, P = 0.066; phase: F[1, 12] = 25.31, P < 0.001; treatment x phase: F[1, 12] = 5.606, P = 0.036) and avoidance (two-way repeated measures ANOVA – treatment: F[1, 12] = 7.25, P = 0.020; phase: F[1, 12] = 19.061, P < 0.001; treatment x phase: F[1, 12] = 7.22, P = 0.020) behavior elicited by the introduction of the aggressor, but not in control animals (**Figure 3CD**). These data further support a selective effect of cortical inhibition on defensive responses to threat.

**Figure 3.**
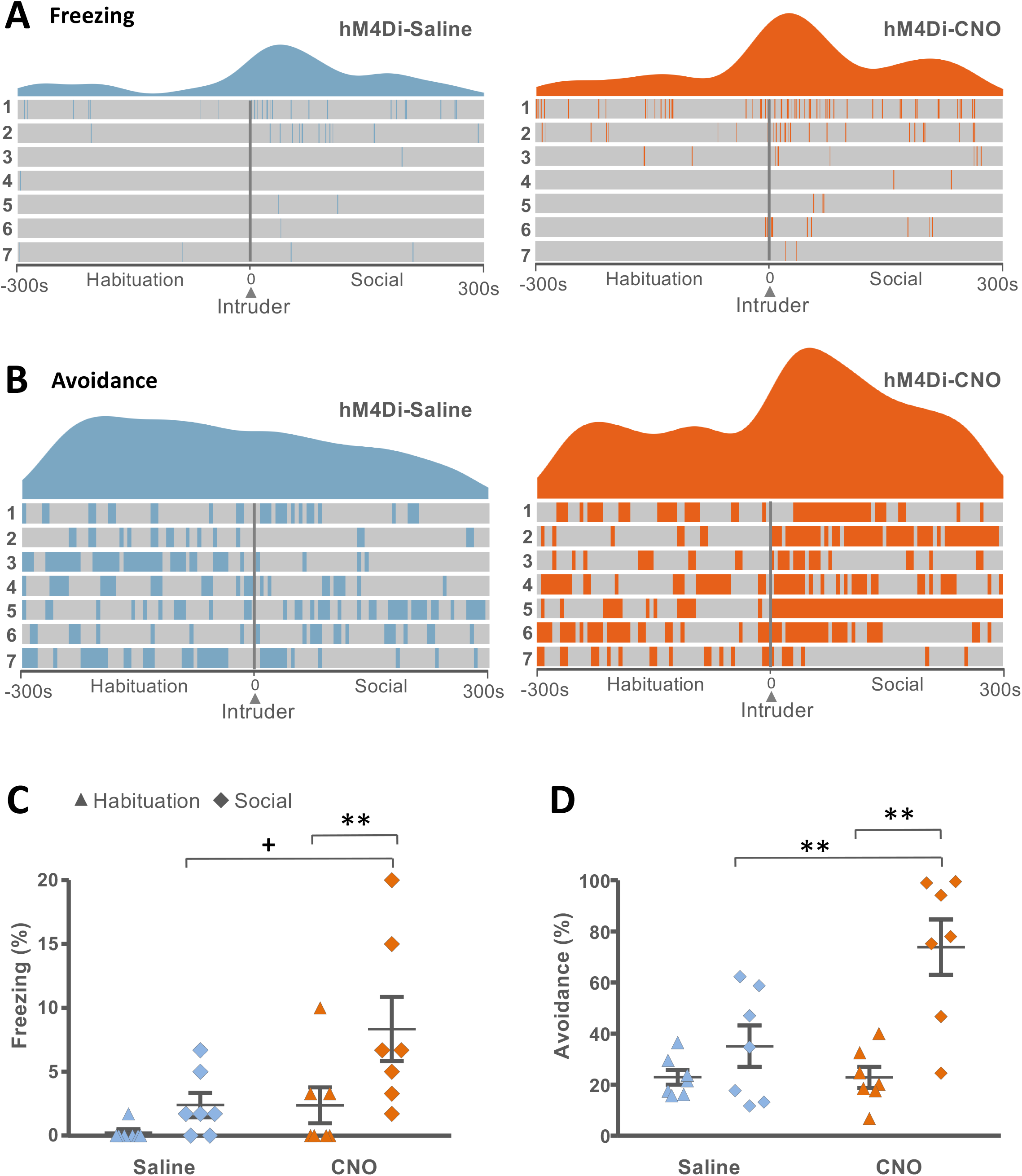
Acute cortical inhibition increases social avoidance and freezing immediately following introduction of the aggressor. Smoothed average and individual data for (**A**) freezing and (**B**) social avoidance of (**left**) saline and (**right**) CNO treated mice during the five minutes before and after introduction of the intruder (arrow). **(C)** Percentage of time spent freezing during one minute before (habituation) and after (social) exposure to the intruder. **(D)** Percentage of time spent in avoidance over one minute before (habituation) and after (social) exposure to the intruder (N = 7; ^+^P < 0.1, *P < 0.05, **P < 0.01, ***P < 0.001).

### Cerebral cortex suppresses prey avoidance

Next, we examined the impact of cortical inhibition on behavioral responses to a novel prey, a powerful non-social threat stimulus. Mice are highly motivated to hunt, kill, and eat cockroaches, but must learn to overcome strong avoidance responses elicited by the prey during the initial encounters (Rossier et al. 2020). To quantify defensive responses to prey, hM4D and control mice were treated with CNO and saline and allowed to habituate to the testing room for 1 hour before the introduction of a live cockroach for forty-five minutes (**Figure 4A**). CNO-treated hM4D mice showed a trend for an increase in defensive freezing (Mann-Whitney: U = 75.0, P = 0.069) and avoidance (Mann-Whitney: U = 76.0, P = 0.079) behavior during the initial phase of exposure to the cockroach when compared to saline-treated hM4D littermates (**Figure 4BC**). However, predatory aggression toward the cockroach was not altered in CNO-treated hM4D mice when compared to saline-treated hM4D mice as shown by an absence of change in time spent attacking, attack duration, latency to kill, and latency to attack (MANOVA – λ = 0.895, F[1, 52] = 1.52, P = 0.209; **Figure 4DE** and **S3**). As expected, CNO treatment did not cause any change in freezing (Mann-Whitney: U=111.0, p= 0.953), avoidance (Mann-Whitney: U=98.0, p= 0.406), or aggressive (MANOVA – λ = 0.895, F[1, 52] = 1.52, P = 0.209) behaviors in control mice (**Figure 4B-E, S3)**. These results support a selective role for cortex in the inhibition of defensive responses to both social and nonsocial threat, but not in the modulation of predatory aggression.

**Figure 4.**
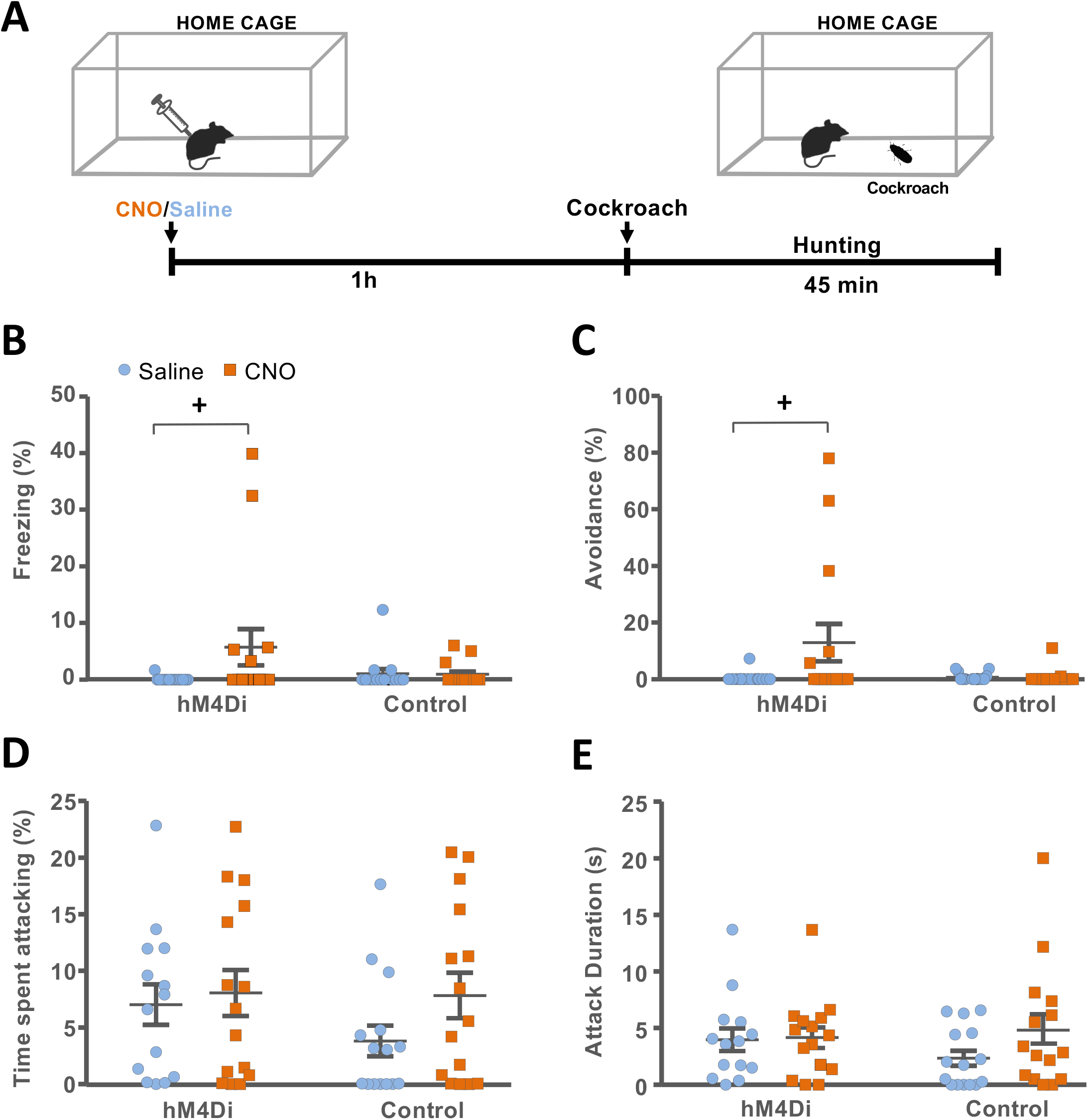
Acute cortical inhibition increases prey avoidance, but not predation. **(A)** Experimental protocol for testing prey avoidance and aggression. Mice were treated with saline or CNO and returned to their home cage for one hour before the introduction of a cockroach and observation for forty-five minutes. Percentage of time spent (**B**) freezing, (**C**) avoiding/cornering, and (**D**) attacking the cockroach and (**E**) mean duration of attacks over the first minute of exposure (N = 14-15; ^+^P < 0.1, *P < 0.05, **P < 0.01, ***P < 0.001).

### Cerebral cortex reduces responses to a physical stressor

Finally, to detect whether cortex has a role in regulating behavioral responses to a non-biological, physical stressor we subjected hM4D and control mice treated with CNO and saline to the forced swim test (FST; Porsolt et al., 1977; Lucki et al., 2001; Can et al., 2012). Mice and rats typically struggle to escape during the initial period of the FST and then settle into a period of alternating escape and immobility as the test continues (Lino-de-Oliveira et al., 2005; Can et al., 2012). The amount of immobility has been used as a measure of stress coping (Molendijk and de Kloet, 2019). Mice were treated with CNO or saline and left to habituate in their home cage for 1 hour before being placed into a beaker containing 3 liters of tepid water where immobility was monitored for four minutes (**Figure 5A**). CNO-treated hM4D mice showed a significant increase in time spent immobile when compared to saline-treated hM4D mice, while the drug treatment had no effect on control animals (two-way ANOVA –genotype: F[1, 54] = 0.307, P = 0.582; treatment: F[1, 54] = 0.819, P = 0.370; genotype x treatment: F[1, 54] = 4.75, P = 0.034; **Figure 5B**).

**Figure 5.**
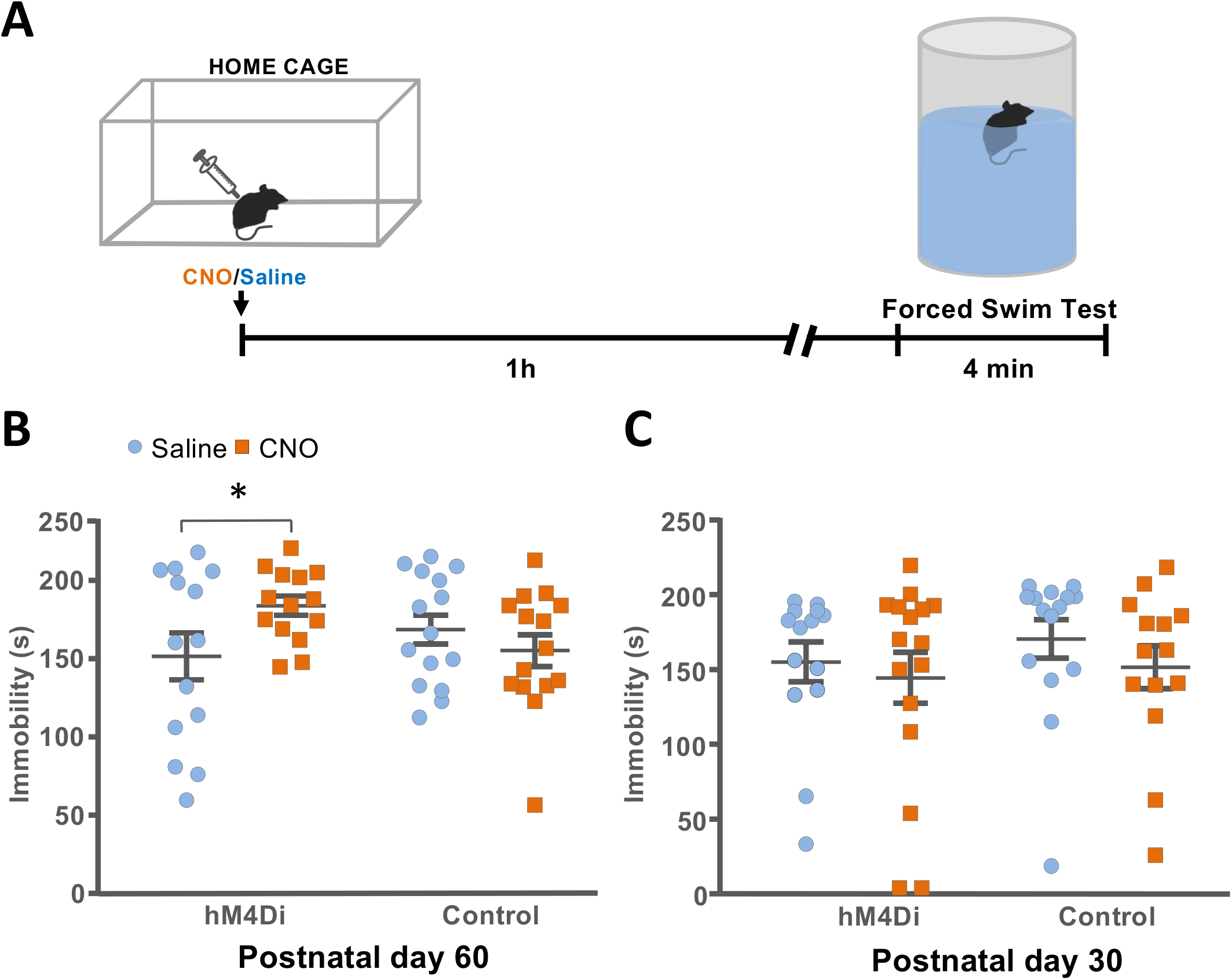
Acute cortical inhibition increases immobility during forced swim stress in adulthood, but not adolescence. **(A)** Experimental protocol for Forced Swim Test. Mice were treated with saline or CNO and returned to their home cage for one hour before introduction into a beaker of water and observation for four minutes. Time spent immobile for (**B**) adult mice (postnatal day 60, P60) and (**C**) adolescent mice (postnatal day 30, P30; N = 14-16; ^+^P < 0.1, *P < 0.05, **P < 0.01, ***P < 0.001).

Based on evidence that decorticated animals show behavioral changes that emerge after sexual maturation (Bjursten et al., 1976) we examined the impact of cortical inhibition on FST responses in adolescent mice (postnatal day 30). At this age, CNO-treated hM4D mice showed no significant change in immobility when compared to saline-treated hM4D littermates (two-way ANOVA – genotype: F[1, 55] = 0.573, P = 0.452; treatment: F[1, 55] = 1.01, P = 0.320; genotype x treatment: F[1, 55] = 0.0793, P = 0.779; **Figure 5C**). CNO treatment had no significant effect in control animals. These results argue for an involvement of cerebral cortex in the inhibition of innate defensive responses to threats regardless of whether they are social, non-social, or physical and that this inhibition may mature during adolescence.

## Discussion

In the present study we examined the behavioral effects of acute pharmacogenetic inhibition of principle neurons in the cerebral cortex of mice. We tested the hypothesis that activity in the cerebral cortex suppresses subcortical structures that promote defensive behaviors and that lesions of cortex are associated with increased defensive responses to a variety of threats. Our findings confirm a selective role for cortex in suppressing innate defensive behaviors, and failed to find any effect of cortical inhibition on other innate behaviors, including predatory aggression. These findings suggest that earlier claims for a disinhibitory effect of cortical lesions on aggression, irritability, and antisocial behavior may be linked to a more specific effect on altering behavioral responses to threat. Our findings provide renewed support for the “release phenomenon” concept described in the eighteenth century (Jackson, 1884) and revisited by scientists in the twentieth century (Macmillan, 1992). This concept argued that the emotional excitement observed in decorticated animals in response to otherwise innocuous stimuli was due to the release of subcortical brain centers from cortical control.

The DREADD-dependent inhibition approach we chose to use allowed us to pharmacologically induce the precise inhibition of principal neurons across the entire neocortex and archicortex and thus overcome confounds associated with the imprecision of earlier aspiration, excitotoxic, or pharmacological lesion studies. Moreover, our pharmacogenetic approach overcomes limitations of earlier irreversible lesions approaches that could have been confounded by long-term compensatory processes. In light of our results in which cortical inhibition showed a selective impact on defensive behaviors, we speculate that either the wider phenotypes described with chronic lesions that frequently included aggression and irritability (Clemente and Lindsley 1967) were the indirect result of a change in threat responses processes, or they were due to compensations arising from the chronic nature of the manipulations.

Our experiment examining the impact of cortical inhibition on prey approach, attack, and capture suggests that cortex modulates prey avoidance, but not aggression. Predatory, or offensive aggression is controlled in major part by distinct neural circuits compared to defensive aggression (Siegel and Pott, 1988; Panksepp, 1992; Miczek et al., 2013) and our findings of normal attack intensity and unaltered latency to attack and kill are consistent with a lack of cortical inhibition on predatory aggression. Similar conclusions have been drawn from cortical lesions performed in cats (De Barenne, 1920; Rothmann, 1923; Cannon and Britton, 1925; Bard, 1928) On the other hand, our social interaction test did not include measurements of social aggression (e.g. resident-intruder aggression) and thus did not allow us to draw conclusions about a role for cortex in defensive aggression.

Cortical inhibition significantly extended active coping responses in the forced swim test (**Figure 5B**) confirming a role for cortical modulation in this test (Duncan et al., 1993; Warden et al., 2012). Although the forced swim test has been widely used as a pharmacological screening method for identifying antidepressants (Lucki et al., 2001), we take a more parsimonious interpretation of our findings to support a role for cortex in the processing of stress-induced defensive escape responses (Commons et al., 2017; Molendijk and de Kloet, 2019). Such an interpretation is consistent with swimming responses in this test reflecting an active coping strategy aimed at escaping from environmental stress (de Kloet and Molendijk, 2016). Consistent with such an interpretation, cortical inhibition increased immobility time (**Figure 5B**) suggesting that cortical activity promotes active escape. However, an alternative interpretation is that the increased swimming we see reflects a reduction of passive coping responses that predominate in the face of inescapable stress (West, 1990).

Importantly, the effect of cortical inhibition was not observed in younger animals (**Figure 5C**). Previous studies have pointed to different mechanism controlling forced swim test behavior in adolescent and mature animals (Doherty et al., 2017) and earlier decortication experiments noted the appearance of behavioral disturbances only after adolescence (Bjursten et al., 1976). We interpret this finding as consistent with the peri-adolescent development of cortical inhibition of behavior reported in humans (Constantinidis and Luna, 2019) and supported by the delayed development of selected cortical-fugal projections in mammals (González-Maeso et al., 2007; Narboux-Nême et al., 2012; Piszczek et al., 2015).

Although our study did not allow us to identify the cortical projects involved in the behavioral effects we observed, our findings are consistent with studies implicating projections from pre-frontal cortex to dorsal periaqueductal grey in the inhibition of social avoidance (Franklin et al., 2017) and to the midbrain dorsal raphe nucleus in the inhibition of forced swim test escape (Warden et al., 2012). Meanwhile, similar inhibitory effects of cortex on brainstem regions involved in defensive behavior have been inferred from human functional connectivity imaging studies. fMRI data in the context of virtual escape from predator threat, demonstrate that the cingulate cortex, hippocampus and amygdala are recruited in the first phase of detection of potential threat and that these regions are subsequently inhibited in concert with the activation of periaqueductal grey when attack is imminent (Mobbs et al., 2007; Mobbs et al., 2009).

Our evoked electrophysiological recordings suggest that the transgenic pharmacogenetic inhibition approach we employed was likely to only partially inhibit cortical excitatory neurotransmission (**Figure 1E**). This finding was not entirely unexpected, as DREADD-based inhibition is frequently found to induce only partial inhibition of both baseline and evoked neural responses when examined in *ex vivo* brain slices. However, we are aware of only one other study in which hM4Di-dependent suppression of neurotransmission was examined *in vivo* (Madroñal et al., 2010). The larger effect seen in this earlier study (60% vs. 25% here) may be related to the moderate expression levels achieved by the transgenic expression of hM4D that, for example, lacks the WPRE RNA stability element (Ray & Dymecki et al., 2011).

It is important to note that the *Emx1*::*Cre* driver line we used is active in both neocortex as well as archicortex. Thus, the full range of cortical regions were suppressed in our experimental animals, including hippocampus, basolateral amygdala, and olfactory cortex (Iwasato et al., 2000; Iwasato et al., 2004). This feature allowed us to systematically target all cortical regions (Kapper, 1909; Garey, 1994; Gerfen and Wilson, 1996; Swanson, 2000) in a manner not possible with earlier physical, chemical, or pharmacological lesions (Bard, 1928; Bard, 1937; Bard and Mountcastle, 1948). However, it leaves open the relative roles of amygdala, hippocampus, and neocortex in the behavioral effects we observe. Notably, lesions of amygdala and hippocampus have been reported to have anxiolytic effects that promote approach behavior (Korn et al., 2017) and abolish innate defensiveness toward a threat (Pentkowski et al., 2006; Martinez et al., 2011), data that goes in the opposite direction of the phenotypes we observed. Thus, we tentatively asign the inhibition of defensive behaviors to neocortex, and speculate that it may be particularly dependent on midline neocortical structures such as prefrontal and cingulate areas that have prominent midbrain projections known to regulate defense (Warden et al., 2012; Franklin et al., 2017; Rozeske et al., 2018).

It remains possible that the increased defensive behavior we observed was a non-specific consequence of a behavioral deficit in our mice. We do not favor such a non-specific effect for several reasons. First, spatial navigation appears not to have been significantly affected by our manipulation despite the known role of neocortical and archicortical structures in this behavior (Poucet et al., 2003; Lopes, 2017; Chersi and Burgess, 2015). For example, we observed no detectable change in hunting behavior (**Figure S3**) or social approach (**Figure S1C**), suggesting the ability of subcortical structures to support sophisticated spatial orientation and goal-directed behaviors. These findings are consistent with earlier cortical lesion studies that found intact navigation, spatial learning, social play, and sexual behavior (Bjursten et al., 1976; Oakley, 1979; Whishaw and Colb 1983; Terry P et al., 1989; Kawai et al., 2015). Second, the persistence of sophisticated social behaviors and their proper habituation across experiences suggests unaffected olfaction. For example, female mice showed no differences in activity, social approach and avoidance toward male intruders (**Figure S1**), while males – whose interactions are known to include territorial threat responses – did (**Figure 2** and **3**). Finally, we observed an increase in escape responses in the forced swim test that does not depend on olfactory cues.

In conclusion, our data point to the existence of a role for cortical structures in suppressing innate defensive responses to a variety of threats. Human functional imaging studies show that prefrontal cortical regions are hyperactivated during active escape from a virtual predator, but quickly shut down when capture by the predator is imminent (Mobbs et al., 2007). This cortical shutdown is accompanied by a hyperactivation of midbrain defense control regions that are known to drive flight behavior and be directly inhibited by prefrontal projections (Franklin et al., 2017). We speculate that this ability may be part of an adaptive mechanism aimed at inhibiting defensive responses under conditions in which cortical circuits can benefit from collecting more information about threat context to guide optimal response strategies, but allow subcortical defensive regions to take over when survival is at stake.

## Materials & Methods

### Animals

All mice tested were obtained by internal colonies from the European Molecular biology laboratory. Mice were maintained in temperature and humidity-controlled condition with food and water provided *ad libitum* and on 12-h light-dark cycle (light on at 7:00). Experimental groups *Emx1*::Cre; *RC*::FPDi (called hM4Di group) and *RC*::FPDi (called Control group) consisted of littermate mice obtained by crossing homozygous *RC*::FPDi mice, kindly provided by Dr. Susan Dymecki (Harvard Medical School, Boston, USA; Ray et al., 2011) and heterozygous *Emx1*::*Cre* mice (Piszczek et al., 2015). Adult CD-1 aggressors used as intruders were selected based on a screening procedure previously described (Franklin et al., 2017). All experiments were performed in accordance with EU Directive 2010/63/EU and under approval of the EMBL Animal Use Committee and Italian Ministry of Health License 541/2015-PR to C.T.G.

### Behavioral testing

#### Social Avoidance Test

Adult male and female animals were singly housed one week before the test day and habituated to saline i.p. injection for two days preceding the test. Experimental mice of the hM4Di or control group were randomly treated with saline or Clozapine-N-oxide (CNO 3mg/kg i.p. in 0.9% saline, Sigma-Aldrich) and placed for 1 hour in their home cage where the test was subsequently carried out (**Figure 2**). After 5 minutes of habituation in which the activity of each subject was automatically recorded an aggressive male CD1 intruder was placed into the resident home cage constrained within a wire-mesh cage. Mice interacted for 5 minutes while approach/avoidance behavior was scored from videotape either automatically in case of activity, social approach, freezing and avoidance or manually for the latency to the first investigation using Solomon coder software. Male and Female behavioral parameters were grouped separately and the test was repeated for five consecutive days.

#### Hunting Test

After treatment with saline or CNO (3mg/kg, i.p.) male and female mice, previously group housed, were isolated for 1 hour in a novel cage where the test occurred. Mice were exposed to a live cockroach (*B. lateralis*). The test ended when the mouse killed the cockroach or after 45 minutes, whichever was shorter. Aggression (time spent attacking, attack duration) was manually scored. Percentage of time spent attacking was calculated as the time spent attacking divided by the total time of the test and the attack duration was calculated as the total time spent attacking divided by the number of attacks. Freezing and avoidance (time spend in the corner of the cage) were quantified during the first minute of exposure to the cockroach.

#### Forced Swim Test

One hour after CNO or saline treatment (3 mg/kg, i.p.) mice (male and female) were placed in a glass cylindrical beaker (26 cm height x 16 cm diameter) filled with 3 liters of tap water at 23-25 C (Can et al., 2012). Rectangular white cardboard dividers where used in order to prevent mice from seeing each other while they were tested side-by-side. The duration of the test was 6 minutes and the first 120 seconds were excluded from behavioral scoring since mice during this period show persistent escape activity (Can et al., 2012). Immobility was calculated by subtracting the time swimming from the total test duration. Swimming was scored whenever the mouse was actively moving its limbs but excluded periods when the animal made only limited circular movements of the legs aimed at maintaining its head above water. After the test, mice were dried with paper towels and placed for a short period under a heat lamp to recover. A subset of mice were tested at both time points, in which case the treatment at P60 was pseudorandomly scrambled.

### In vivo *electrophysiology*

Synaptic field potentials (fEPSPs) were evoked by delivering 100 μs, square, biphasic pulses applied to the contralateral motor cortex (0.1 Hz, 40-50% of maximum response amplitude, 4 minutes) at 40, 20 and 5 minutes before and 20, 40, 60 and 80 min after drug treatment using a pulse generator (CS-420, Cibertec) connected to an electrical stimulator (ISU-200bip, Cibertec). The neural signal was amplified (gain 1000x) and filtered (bandwidth 0.1-3 kHz) through a headstage 20x preamplifier and a 50x differential amplifier (Omniplex, Plexon). Signals were digitized at 1600 Hz and continuous recordings were collected for post hoc analysis. For initial studies the peak-to-peak fEPSP amplitude values were extracted using commercial software (Spike2 and SIGAVG, Cambridge Electronic Design)

Electrode placement was confirmed by eliciting an electrolytic lesion at the recording site (motor cortex) and the mice were anesthetized three days later using 2.5% Avertin (400 mg/kg, i.p.; Sigma-Aldrich) and perfused transcardially (4.0% vol paraformaldehyde, 0.1M phosphate buffer, pH 7.4). Serial coronal sections (50 microns) were collected using a vibratome (Leica VT1000S) and mounted with DAPI in the mounting medium (MOWIOL; 1:1000). Sections were visualized with a fluorescent microscope (Leica DFC 345 FX) and the green filter was used to visualize the autofluorescence generated by the damage created in the tissue by the electrolytic lesion.

### Histology

Mice were deeply anesthetized with Avertin (400 mg/kg, i.p.; Sigma-Aldrich), perfused transcardially (4% paraformaldehyde in 0.1 M phosphate buffer, pH 7.4) and brains were post-fixed in 4% paraformaldehyde. Brains were cryoprotected and incubated overnight in a solution containing 30% sucrose in 0.1 M PBS and then frozen in Tissue-Teck OCT compound. Coronal slices (50 microns) were cut with a sliding cryostat, mounted on SuperPlus slides, allowed to dry at 42 C on a flattening table (Leica HI1220) and boiled in citrate buffer (10 mM). Sections were incubated with a blocking solution (1% BSA, 5% NGS in PBS, 0.4% Triton X-100) for 1 hour. A primary antibody was used to detect HA (anti-rat, 1:200, 11867423001, Roche) and detection carried out with fluorescent-labeled secondary antibodies (goat anti-rat, 1:800, Alexa Fluor 488, A-101650, Invitrogen). For the detection of mCherry endogenous fluorescent protein animals were anesthetized with Avertin, transcardically perfused and brains post-fixed overnight in 4% paraformaldehyde. Coronal slices (50 microns) were cut with a sliding cryostat (Leica) and mCherry was imaged with a fluorescent microscope (Leica DFC 345 FX, 10X/0.3, 40x/0.8).

### Statistical analysis

All data analysis was performed using Sigmaplot software and all data are reported as mean ± s.e.m. Statistical significance was determined by two-way ANOVA followed by Tukey *post hoc* for Activity, Social Approach and Forced Swim Test measures. For Latency to Investigate in the social approach test and Freezing and Avoidance in the hunting test non-parametric Mann-Whitney U-tests was used. Two-way repeated measure ANOVA followed by *post hoc* Bonferroni testing was used to determine the significance for freezing and avoidance in the Social Approach test and twoway repeated measures ANOVA followed by *post hoc* Student-Newman-Keuls testing for *in vivo* electrophysiology experiments. Time spent attacking, attack duration, latency to kill, and latency to attack were analyzed by multiple analysis of variance (MANOVA).

## Author contributions

All behavioral experiments and data analysis were carried out by S.N.; M.E.M. and S.N. carried out electrophysiological experiments and together with C.T.G. designed the experiment; S.N. together with S.D. carried out the Forced Swim Test; S.N., S.D., and C.T.G. designed the experiments, C.T.G supervised the project and together with S.N. conceived the project and wrote the manuscript.

## Acknowledgments

We thank Daniel Rossier for support with hunting behavior, Roberto Voci and Valerio Rossi for mouse husbandry, and the EMBL Microscopy and Laboratory Animal Resources Facilities. The work was supported by EMBL and the European Research Council (ERC) Advanced Grant COREFEAR to C.T.G.

## Supplementary Figure Legends

**Figure S1.**
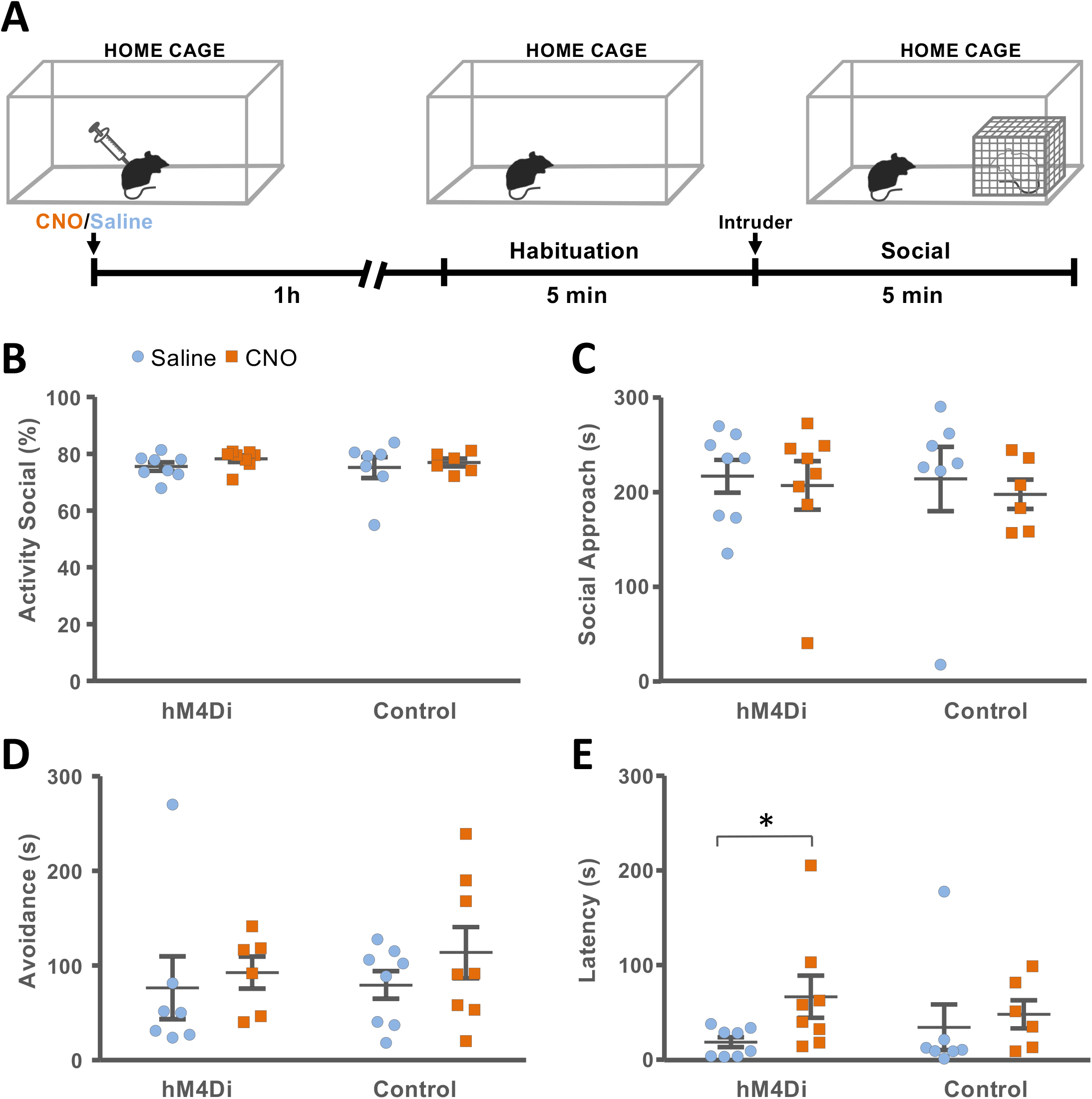
Acute cortical inhibition fails to alter social avoidance in female mice. (**A**) Experimental protocol for testing social avoidance. Mice were treated with saline or CNO and returned to their home cage for one hour before baseline testing for five minutes followed by the introduction at one side of the home cage of an aggressor confined to a wire mesh cage and further observation for five minutes. (**B**) Activity of mice measured during the social test phase, (**C**) social approach, (**D**) social avoidance, and (**E**) latency to first social approach (N = 6-8, ^+^P < 0.1, *P < 0.05, **P < 0.01, ***P < 0.001).

**Figure S2.**
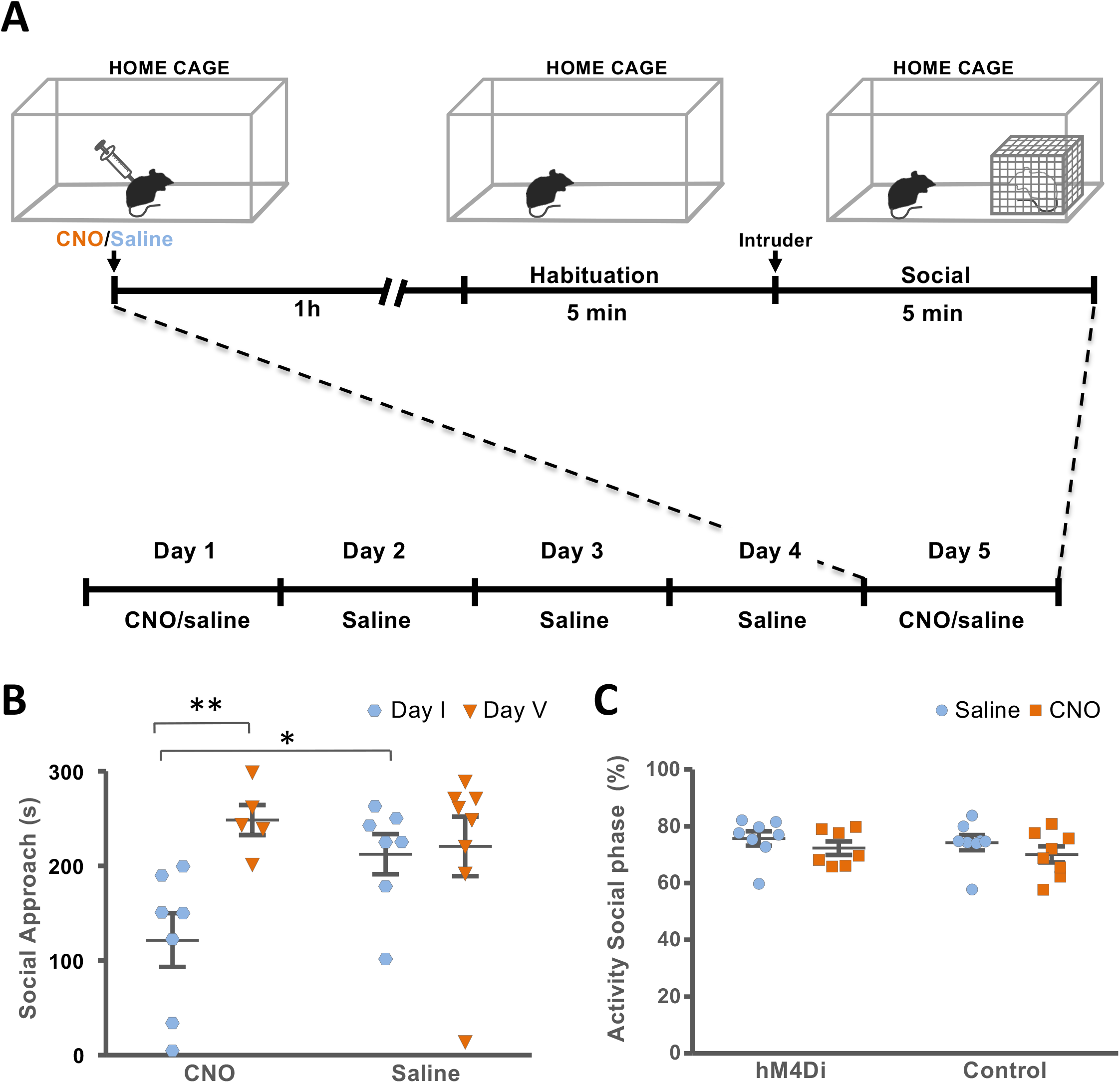
Acute cortical inhibition fails to alter social avoidance after threat habituation. (**A**) Social avoidance testing was carried out each day for five consecutive days as above. Mice were treated with saline or CNO on the first and last day and with saline on all other days. (**B**) Social Approach on the first and last day of testing for mice in the hM4Di experimental group. **(C**) Activity during social phase for mice on the last day of testing (N = 5-8; ^++^P < 0.1, *P < 0.05, **P < 0.01, ***P < 0.001).

**Figure S3.**
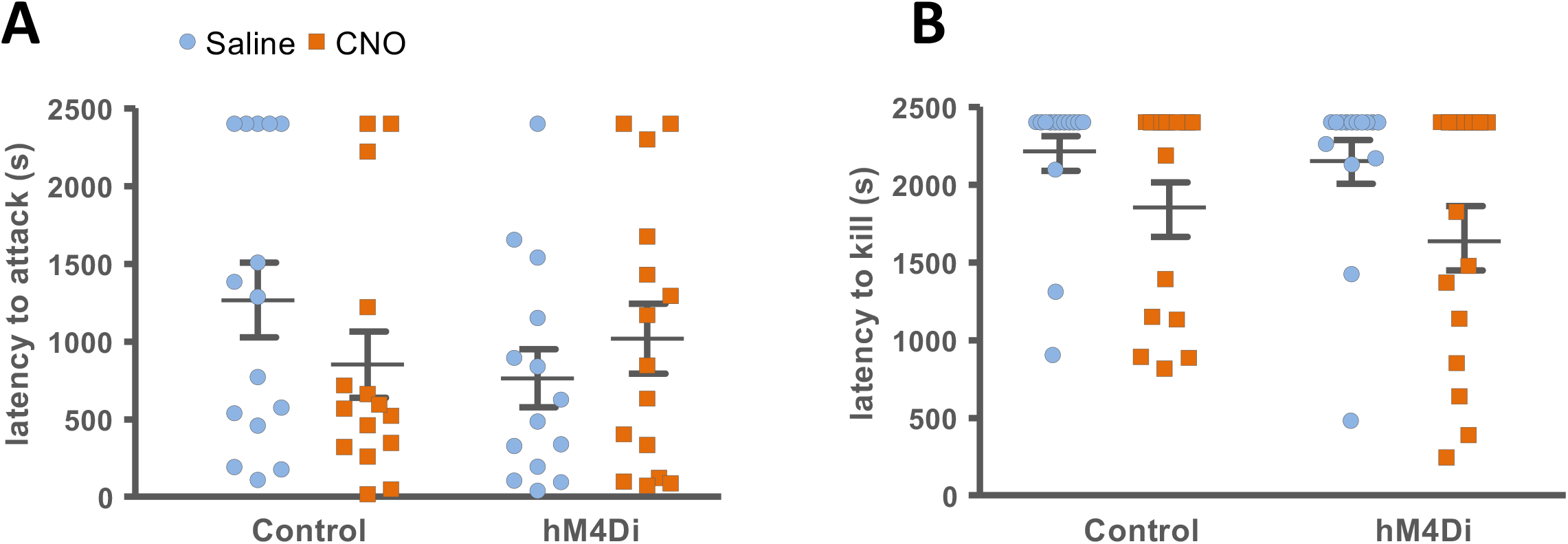
Acute cortical inhibition fails to alter predatory aggression. Latency to (**A**) attack and **(B)** kill prey in the predatory aggression assay (N = 14-15).

## Notes

### Competing Interest Statement

The authors have declared no competing interest.

## References

Adams DB (1979) Brain mechanisms for offense, defense, and submission. Behavioral and Brain Sciences 2:201–213

Armbruster BN, Li X, Pausch MH, Herlitze S, Roth BL (2007) Evolving the lock to fit the key to create a family of G protein-coupled receptors potently activated by an inert ligand. Proc Natl Acad Sci U S A 104:5163–5168.

Bard P, Mountcastle V B (1948) Some forebrain mechanisms involved in expression of rage with special reference to suppression of angry behavior. Association for Research in Nervous and Mental Disease 27:362–404

Bard P (1928) A diencephalic mechanism for the expression of rage with special reference to the sympathetic nervous system American Journal of Physiology 84:490–515.

Bard P (1934) On emotional expression after decortication with some remarks on certain theoretical views: Part II. Psychological Review 41:424–449

Bard P, Rioch DM (1937) A study of four cats deprived of neocortex and additional portions of the forebrain. Johns Hopkins Medical Journal 60:73–147.

Bernhard CG, Bohm E (1954) Cortical representation and functional significance of the corticomotoneuronal system. AMA Arch Neurol Psychiatry 72:473–502.

Bjursten LM, Norrsell K, Norrsell U (1976) Behavioural repertory of cats without cerebral cortex from infancy. Exp Brain Res 25:115–130.

Can A, Dao DT, Arad M, Terrillion CE, Piantadosi SC, Gould TD (2012) The mouse forced swim test. J Vis Exp:e3638.

Cannon WB and Britton SW (1925) Pseudoaffective medulliadrenal secretion. American Journal of Psychology 72:283–294.

Clemente C D, Lindsley DB (1967) Aggression and Defense: Neural Mechanisms and Social Patterns. Brain Function Vol. 1

Chersi F, Burgess N (2015) The Cognitive Architecture of Spatial Navigation: Hippocampal and Striatal Contributions. Neuron 88:64–77.

Commons KG, Cholanians AB, Babb JA, Ehlinger DG (2017) The Rodent Forced Swim Test Measures Stress-Coping Strategy, Not Depression-like Behavior. ACS Chem Neurosci 8:955–960.

Constantinidis C, Luna B (2019) Neural Substrates of Inhibitory Control Maturation in Adolescence. Trends Neurosci 42:604–616.

de Kloet ER, Molendijk ML (2016) Coping with the Forced Swim Stressor: Towards Understanding an Adaptive Mechanism. Neural Plast 2016:6503162.

Dalgleish T, Dunn BD, Mobbs D (2009) Affective Neuroscience: Past, Present, and Future. Emotion Review 1:355–368

De Barenne D, J. G. (1920) Recherches Expérimentales sur le Functions du Sistème Nervaux central Faites en Particulier sur deux chats dont le néopallium evait été enlevé. Arch. Néerl. de Physiol 4:31–123

Doherty TS, Blaze J, Keller SM, Roth TL (2017) Phenotypic outcomes in adolescence and adulthood in the scarcity-adversity model of low nesting resources outside the home cage. Dev Psychobiol 59:703–714.

Duncan GE, Johnson KB, Breese GR (1993) Topographic patterns of brain activity in response to swim stress: assessment by 2-deoxyglucose uptake and expression of Fos-like immunoreactivity. J Neurosci 13:3932–3943.

Franklin TB, Silva BA, Perova Z, Marrone L, Masferrer ME, Zhan Y, Kaplan A, Greetham L, Verrechia V, Halman A, Pagella S, Vyssotski AL, Illarionova A, Grinevich V, Branco T, Gross CT (2017) Prefrontal cortical control of a brainstem social behavior circuit. Nat Neurosci 20:260–270.

Garey LJ (1994) Brodmann’s‘Localisation in the Cerebral Cortex’, London:Smith-Gordon.

Gerfen CR, Wilson CJ (1996) The basal ganglia, in: L.W. Swanson, A. Bjoörklund, T. Hoökfelt (Eds.), Handbook of Chemical Neuroanatomy, Integrated Systems of the CNS, Part III 12:371–468.

Goltz F (1881) Discussion of the localization of function in the cortex cerebri [in German]. Transactions of the International Medical Congress 1:218–228.

Goltz F (1888) Uber die verrichtungen des grosshirns. Pfugers Arch 42:419–467.

Goltz F (1892) Der hund ohne großhirn. Pflüg Arch ges Physiol 51:570–614.

González-Maeso J, Weisstaub NV, Zhou M, Chan P, Ivic L, Ang R, Lira A, Bradley-Moore M, Ge Y, Zhou Q, Sealfon SC, Gingrich JA (2007) Hallucinogens recruit specific cortical 5-HT(2A) receptor-mediated signaling pathways to affect behavior. Neuron 53:439–452.

Grenier F, Timofeev I, Steriade M (1998) Leading role of thalamic over cortical neurons during postinhibitory rebound excitation. Proc Natl Acad Sci U S A 95:13929–13934.

Harlow JM (1869) Recovery from the passage of an iron bar through the head. Publications of the Massachusetts Medical Society 2:327–347.

Head H (1921) Croonian Lecture: Release of Function in the Nervous System. Proceeding of the royal society 92:184–209

Heindorf M, Arber S, Keller GB (2018) Mouse Motor Cortex Coordinates the Behavioral Response to Unpredicted Sensory Feedback. Neuron 99:1040–1054.e1045.

Iwasato T, Nomura R, Ando R, Ikeda T, Tanaka M, Itohara S (2004) Dorsal telencephalon-specific expression of Cre recombinase in PAC transgenic mice. Genesis 38:130–138.

Iwasato T, Datwani A, Wolf AM, Nishiyama H, Taguchi Y, Tonegawa S, Knöpfel T, Erzurumlu RS, Itohara S (2000) Cortex-restricted disruption of NMDAR1 impairs neuronal patterns in the barrel cortex. Nature 406:726–731.

Jackson JH (1884) The Croonian Lectures on Evolution and Dissolution of the Nervous System. Br Med J 1:703–707.

Kapper CUA (1909) The phylogenesis of the palaeo-cortex and archi-cortex compared with the evolution of the visual neo-cortex, Arch. Neurol. Psychiat 4:161–173.

Kawai R, Markman T, Poddar R, Ko R, Fantana AL, Dhawale AK, Kampff AR, Ölveczky BP (2015) Motor cortex is required for learning but not for executing a motor skill. Neuron 86:800–812.

Korn CW, Vunder J, Miró J, Fuentemilla L, Hurlemann R, Bach DR (2017) Amygdala Lesions Reduce Anxiety-like Behavior in a Human Benzodiazepine-Sensitive Approach-Avoidance Conflict Test. Biol Psychiatry 82:522–531.

Lawrence DG, Kuypers HG (1968) The functional organization of the motor system in the monkey. I. The effects of bilateral pyramidal lesions. Brain 91:1–14.

Lopes G, Nogueira J, Dimitriadis G, Menendez JA, Paton JJ, Kampff AR (2017) A robust role for motor cortex. bioRxix

Lino-de-Oliveira C, De Lima TC, de Pádua Carobrez A (2005) Structure of the rat behaviour in the forced swimming test. Behav Brain Res 158:243–250.

Lucki I, Dalvi A, Mayorga AJ (2001) Sensitivity to the effects of pharmacologically selective antidepressants in different strains of mice. Psychopharmacology (Berl) 155:315–322.

Macmillan M (1992) Inhibition and the control of behavior.From Gall to Freud via Phineas Gage and the frontal lobes. Brain Cogn 19:72–104.

Madroñal N, Delgado-García JM, Fernández-Guizán A, Chatterjee J, Köhn M, Mattucci C, Jain A, Tsetsenis T, Illarionova A, Grinevich V, Gross CT, Gruart A (2016) Rapid erasure of hippocampal memory following inhibition of dentate gyrus granule cells. Nat Commun 7:10923.

Marple-Horvat DE, Amos AJ, Armstrong DM, Criado JM (1993) Changes in the discharge patterns of cat motor cortex neurones during unexpected perturbations of on-going locomotion. J Physiol 462:87–113.

Martinez RC, Carvalho-Netto EF, Ribeiro-Barbosa ER, Baldo MV, Canteras NS (2011) Amygdalar roles during exposure to a live predator and to a predator-associated context. Neuroscience 172:314–328.

Merel J, Botvinick M, Wayne G (2019) Hierarchical motor control in mammals and machines. Nat Commun 10:5489.

Miczek KA, de Boer SF, Haller J (2013) Excessive aggression as model of violence: a critical evaluation of current preclinical methods. Psychopharmacology (Berl) 226:445–458.

Mizutani H, Hori T, Takahashi T (2006) 5-HT1B receptor-mediated presynaptic inhibition at the calyx of Held of immature rats. European Journal of Neuroscience 24:1946–54

Mobbs D, Petrovic P, Marchant JL, Hassabis D, Weiskopf N, Seymour B, Dolan RJ, Frith CD (2007) When fear is near: threat imminence elicits prefrontal-periaqueductal gray shifts in humans. Science 317:1079–1083.

Mobbs D, Marchant JL, Hassabis D, Seymour B, Tan G, Gray M, Petrovic P, Dolan RJ, Frith CD (2009) From threat to fear: the neural organization of defensive fear systems in humans. J Neurosci 29:12236–12243.

Molendijk ML, de Kloet ER (2019) Coping with the forced swim stressor: Current state-of-the-art. Behav Brain Res 364:1–10.

Molnár Z, Kaas JH, de Carlos JA, Hevner RF, Lein E, Němec P (2014) Evolution and development of the mammalian cerebral cortex. Brain Behav Evol 83:126–139.

Montiel JF, Aboitiz F (2015) Pallial patterning and the origin of the isocortex. Front Neurosci 9:377.

Narboux-Nême N, Evrard A, Ferezou I, Erzurumlu RS, Kaeser PS, Lainé J, Rossier J, Ropert N, Südhof TC, Gaspar P (2012) Neurotransmitter release at the thalamocortical synapse instructs barrel formation but not axon patterning in the somatosensory cortex. J Neurosci 32:6183–6196.

Oakley DA (1979) Learning with food reward and shock avoidance in neodecorticate rats. Exp Neurol 63:627–642.

Oh SW et al. (2014) A mesoscale connectome of the mouse brain. Nature 508:207–214.

Panksepp J (1992) A critical role for “affective neuroscience” in resolving what is basic about basic emotions. Psychol Rev 99:554–560.

Pentkowski NS, Blanchard DC, Lever C, Litvin Y, Blanchard RJ (2006) Effects of lesions to the dorsal and ventral hippocampus on defensive behaviors in rats. Eur J Neurosci 23:2185–2196.

Piszczek L, Piszczek A, Kuczmanska J, Audero E, Gross CT (2015) Modulation of anxiety by cortical serotonin 1A receptors. Front Behav Neurosci 9:48.

Porsolt RD, Le Pichon M, Jalfre M (1977) Depression: a new animal model sensitive to antidepressant treatments. Nature 266:730–732.

Poucet B, Lenck-Santini PP, Paz-Villagrán V, Save E (2003) Place cells, neocortex and spatial navigation: a short review. J Physiol Paris 97:537–546.

Ray RS, Corcoran AE, Brust RD, Kim JC, Richerson GB, Nattie E, Dymecki SM (2011) Impaired respiratory and body temperature control upon acute serotonergic neuron inhibition. Science 333:637–642.

Rossier D, La Franca V, Salemi T, Gross CT (2020) A neural circuit for competing approach and avoidance underlying prey capture. BioRxiv

Rothmann H (1923). Zusammenfassender bericht über den rothmannschen gehirnlosen hund nach klinisher hund anatomischer untersuchung. Z. ges. Neurol. Psychiat. 87:247–313

Rozeske RR, Jercog D, Karalis N, Chaudun F, Khoder S, Girard D, Winke N, Herry C (2018) Prefrontal-Periaqueductal Gray-Projecting Neurons Mediate Context Fear Discrimination. Neuron 97:898–910.e896.

Schröder P, Schmidt TT, Blankenburg F (2019) Neural basis of somatosensory target detection independent of uncertainty, relevance, and reports. Elife 8.

Siegel A, Pott CB (1988) Neural substrates of aggression and flight in the cat. Prog Neurobiol 31:261–283.

Spain WJ, Schwindt PC, Crill WE (1991) Post-inhibitory excitation and inhibition in layer V pyramidal neurones from cat sensorimotor cortex. J Physiol 434:609–626.

Swanson LW (2000) Cerebral hemisphere regulation of motivated behavior. Brain Res 886:113–164.

Schaltenbrand G, Cobb S (1930) Clinical and anatomical studies on two cats without neocoetex. Brain: A Journal of Neurology 53:449–488.

Terry P, Herbert BA, Oakley DA. Anomalous patterns of response learning and transfer in decorticate rats. 1989 Behavioural brain research 33:105–109

Tessmar-Raible K, Raible F, Christodoulou F, Guy K, Rembold M, Hausen H, Arendt D (2007) Conserved sensory-neurosecretory cell types in annelid and fish forebrain: insights into hypothalamus evolution. Cell 129:1389–1400.

Tomer R, Denes AS, Tessmar-Raible K, Arendt D (2010) Profiling by image registration reveals common origin of annelid mushroom bodies and vertebrate pallium. Cell 142:800–809.

van Bergen RS, Ma WJ, Pratte MS, Jehee JF (2015) Sensory uncertainty decoded from visual cortex predicts behavior. Nat Neurosci 18:1728–1730.

Warden MR, Selimbeyoglu A, Mirzabekov JJ, Lo M, Thompson KR, Kim SY, Adhikari A, Tye KM, Frank LM, Deisseroth K (2012) A prefrontal cortex-brainstem neuronal projection that controls response to behavioural challenge. Nature 492:428–432.

West AP (1990) Neurobehavioral studies of forced swimming: the role of learning and memory in the forced swim test. Prog Neuropsychopharmacol Biol Psychiatry 14:863–877.

Winans SS (1967) Visual form discrimination after removal of the visual cortex in cats. Science 158:944–946.

Whishaw IQ, Kolb B (1983) Can Male Decorticate Rats Copulate? Behavioral Neuroscience 97:270–279

Yang HW, Lemon RN (2003) An electron microscopic examination of the corticospinal projection to the cervical spinal cord in the rat: lack of evidence for cortico-motoneuronal synapses. Exp Brain Res 149:458–469.

